# Interpreting Wide-Band Neural Activity Using Convolutional Neural Networks

**DOI:** 10.1101/871848

**Authors:** Markus Frey, Sander Tanni, Catherine Perrodin, Alice O’Leary, Matthias Nau, Jack Kelly, Andrea Banino, Daniel Bendor, Christian F. Doeller, Caswell Barry

**Author notes:** Joined senior authors (Christian F. Doeller, Caswell Barry).

## Abstract

Rapid progress in technologies such as calcium imaging and electrophysiology has seen a dramatic increase in the size and extent of neural recordings. Even so, interpretation of this data often depends on manual operations and requires considerable knowledge about the nature of the representation. Decoding provides a means to infer the information content of such recordings but typically requires highly processed data and prior knowledge of the encoding scheme. Here, we developed a deep-learning-framework able to decode sensory and behavioural variables directly from wide-band neural data. The network requires little user input and generalizes across stimuli, behaviours, brain regions, and recording techniques. Once trained, it can be analysed to determine elements of the neural code that are informative about a given variable. We validated this approach using data from rodent auditory cortex and hippocampus, identifying a novel representation of head direction encoded by putative CA1 interneurons.

## Introduction

A central aim of neuroscience is deciphering the neural code, understanding the neural representation of sensory features and behaviours, as well as the computations that link them. The task is complex, and although there have been notable successes - such as the identification of orientation selectivity in V1 (Hubel and Wiesel, 1959) and the representation of self-location provided by hippocampal place cells (O’Keefe and Dostrovsky, 1971) - progress has been slow. Neural activity is high dimensional and often sparse, while the available datasets are typically incomplete, being both temporally and spatially limited. This problem is compounded by the fact that the code is multiplexed and functionally distributed (Walker et al., 2011). As such, activity in a single region may simultaneously represent multiple variables, to differing extents, across different elements of the neural population. Taking the entorhinal cortex for example, a typical electrophysiological recording might contain spike trains from distinct cells predominantly encoding head direction, self-location, and movement speed via their firing rates (Sargolini et al., 2006; Kropff et al., 2015; Hafting et al., 2005), while other neurons have more complex composite representations (Hardcastle et al., 2017). At the same time, information about speed and location can also be identified from the local field potential (LFP) (McFarland et al., 1975) and the relative timing of action potentials (O’Keefe and Recce, 1993). Fundamentally, although behavioural states and sensory stimuli can generally be considered to be low dimensional, finding the mapping between noisy neural representations and these less complex phenomena is far from trivial.

Historically, the approach for identifying the correspondence between neural data and external observable states – stimuli or behaviour – has been one of raw discovery. An experimenter, guided by existing knowledge, must recognise the fact that the activity covaries with some other factor. Necessarily this is an incremental process, favouring identification of the simplest and most robust representations, such as the sparse firing fields of place cells (Muller et al., 1987). Classical methods, like linear regression and linear-nonlinear-Poisson cascade models (Corrado et al., 2005; Kropff et al., 2015), provide powerful tools for the characterisation of existing representations but are less useful for the identification of novel responses - they typically require highly processed data in conjunction with strong assumptions about the neural response, and in the former cases are limited to one dimensional variables. Recent advances in machine learning suggest an alternative strategy. Artificial neural networks (ANNs) trained using error backpropagation regularly exceed human-level performance on tasks in which high dimensional data is mapped to lower dimensional labels (Krizhevsky et al., 2012; Mnih et al., 2015). Indeed, these tools have successfully been applied to *processed* neural data - accurately decoding behavioural variables from observed neural firing rates (Glaser et al., 2017; Tampuu et al., 2018). However, the true advantage of ANNs is not their impressive accuracy but rather the fact that they make few assumptions about the structure of the input data and, once trained, can be analysed to determine which elements of the input, or indeed combination of elements, are most informative (Cichy and Kaiser, 2019; Hasson et al., 2020; Cammarata et al., 2020). Moreover, this framework provides full control over the weights, activations, and objective functions of the model, allowing fine-grained analysis of the inner workings of the network. Viewed in this way ANNs potentially provide a means to accelerate the discovery of novel neural representations.

To test this proposal, we developed a convolutional network (LeCun et al., 2015) able to take minimally processed, wide-band neural data as input and output predicted continuous regression variables. In the first instance, we trained the model with unfiltered and unclustered electrophysiological recordings made from the CA1 pyramidal cell layer in freely foraging rodents. As expected, the network accurately decoded the animals’ location, speed, and head direction - without spike sorting or additional user input. Analysis of the trained network showed that it had ‘discovered’ place cells (O’Keefe and Dostrovsky, 1971; O’keefe and Nadel, 1978) - frequency bands associated with pyramidal waveforms being highly informative about self-location (Epsztein et al., 2011). Equally, it successfully recognized that theta-band oscillations in the LFP were informative about running speed (McFarland et al., 1975; Jeewajee et al., 2008). Unexpectedly, the network also identified a population of putative CA1 interneurons that encoded information about head direction. We corroborated this observation using conventional tools, confirming that the firing rate of these neurons was modulated by facing-direction, a previously unreported relationship. Beyond this we found the trained network provided a means to efficiently conduct analyses which would otherwise have been complex or time consuming. For example, comparison of all frequency bands revealed positive interactions between frequencies associated with waveforms - components of the neural code that convey more information together than when considered individually. Subsequently, to demonstrate the generality of this approach, we applied the same architecture to electrophysiological data from auditory cortex as well as two-photon calcium imaging data (Stosiek et al., 2003) acquired while mice explored a virtual environment.

Our model differs markedly from conventional decoding methods which typically use Bayesian estimators (Zhang et al., 1998) in conjunction with highly processed neural data. In the case of extracellular recordings, this usually implies that time-series are filtered and processed to detect action potentials and assign them to specific neurons. Necessarily this discards information in frequency bands outside of the spike range, potentially introducing biases implicit in the algorithm used (Pachitariu et al., 2016; Chung et al., 2017; Lee et al., 2017) and operator’s subjective preferences (Harris et al., 2000; Wood et al., 2004), and - despite considerable advances - still demands considerable manual input to adjust clusters (Pachitariu et al., 2016). Furthermore, accurate calculation of prior expectations regarding the way in which the data varies with the decoded variable – an essential component of Bayesian decoding – requires considerable knowledge about the structure of the neural signal being studied and appropriate noise models. Other authors have attempted to address some of these shortcomings, for example, decoding without assigning action potentials to specific neurons (Kloosterman et al., 2013; Ackermann et al., 2019; Deng et al., 2015) or combining LFP and spiking data (Stavisky et al., 2015) for cursor control in patients. However, these approaches did not use wide-band unprocessed data and relied on existing assumptions about neural coding statistics, while their primary focus was simply to improve decoding accuracy. In contrast, the flexible, general-purpose approach we describe here requires few assumptions and - once trained - can be interrogated to inform the discovery of novel neural representations. In addition, as the model does not rely on specific oscillations or spike waveforms, it can easily generalize across domains - a fact we demonstrate with optical imaging data.

## Results

### Accurate decoding of self-location from CA1 recordings

In the first instance we sought to evaluate our network-based decoding approach on well characterised neural data with a clear behavioural correlate. To this end we used as input extracellular electrophysiological signals recorded from hippocampal region CA1 in five freely moving rats - place cells from this area being noted for their spatially constrained firing fields that convey considerable information about an animals self-location (O’Keefe and Dostrovsky, 1971; Muller et al., 1987). Animals were bilaterally implanted with 32 tetrodes and, after recovery and screening, 128 channel wide-band (0Hz to 15000 Hz sampled at 30kHz) recordings were made while the rats foraged in a 1.25 × 1.75m arena for approximately 40 minutes (see methods). Raw electrophysiological data were decomposed using Morlet wavelets to generate a three-dimensional representation depicting time, channels, and frequencies from 2Hz to 15000Hz (Figure 1A) (Torrence and Compo, 1998). Using the wavelet coefficients as inputs, the model was trained in a supervised-fashion using error backpropagation with the X and Y coordinates of the animal as regression targets.

**Figure 1:**
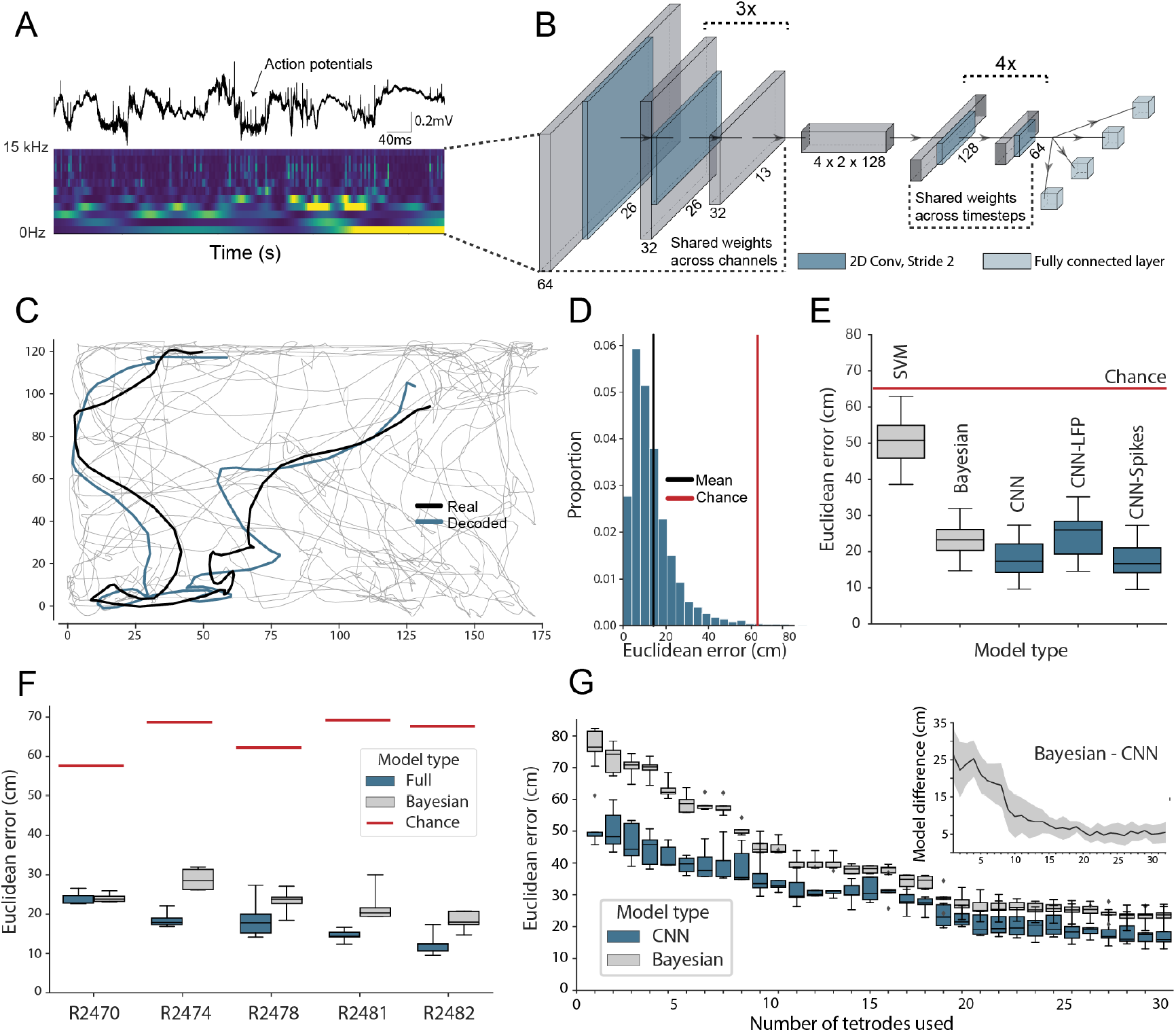
Accurate decoding of self-location from unprocessed hippocampal recordings. A) Top, a typical ‘raw’ extracellular recording from a single CA1 electrode. Bottom, wavelet decomposition of the same data, power shown for frequency bands from 2Hz to 15kHz (bottom to top row). B) At each timestep wavelet coefficients (64 time points, 26 frequency bands, 128 channels) were fed to a deep network consisting of 2D convolutional layers with shared weights, followed by a fully-connected layer with a regression head to decode self-location; schematic of architecture shown. C) Example trajectory from R2478, true position (black) and decoded position (blue) shown for 3s of data. Full test-set shown in **Video 1**. D) Distribution of decoding errors from trial shown in (C), mean error (14.2cm ± 12.9cm, black), chance decoding of self-location from shuffled data (62.2cm ± 9.09cm, red). E) Across all five rats, the network (CNN) was more accurate than a machine learning baseline (SVM) and a Bayesian decoder (Bayesian) trained on action potentials. This was also true when the network was limited to high frequency components (>250Hz, CNN-Spikes). When only local frequencies were used (<250Hz, CNN-LFP), network performance dropped to the level of the Bayesian decoder (distributions show the five-fold cross validated performance across each of five animals, n=25). F) Decoding accuracy for individual animals, the network outperformed the Bayesian decoder in all cases. An overview of the performance of all tested models can be seen in Figure S2. G) The advantage of the network over the Bayesian decoder increased when the available data was reduced by downsampling the number of channels (data from R2478). Inset shows the difference between the two methods.

To reduce computational load and improve test set generalisation we use 2D-convolutions with shared weights applied to the three-dimensional input (Figure 1B, Table S1) - the first eight convolutional layers having weights shared across channels and the final six across time. Implementing weight sharing in this way is desirable as the model is able to efficiently identify features that reoccur across time and channels, for example, prominent oscillations or waveforms, while also drastically reducing model complexity. For comparison, an equivalent architecture trained to decode position from 128 channels of hippocampal electrophysiological but without shared weights had 38,144,900 hyperparameters compared to 5,299,653 - an increase of 720%. The more complex model took 4.7 hours to run per epoch, as opposed to 175s, and ultimately yielded less accurate decoding (Figure S1).

The model accurately decoded position from the unprocessed neural data in all rats, providing a continuous estimate of location with an average error less than 10% of the environment’s length. This demonstrates that, as expected, the network was able to identify informative signals in the raw neural data (Mean error 17.31cm ± 4.46cm; Median error 11.40cm ± 3.82cm; Chance level 65.03cm ± 6.91cm; Figure 1C,D). To provide a familiar benchmark, we applied a standard Bayesian decoder with a continuity prior (Zhang et al., 1998; Ólafsdóttir et al., 2015) to the spiking data from the same datasets (see methods). To this end, action potentials were identified, clustered, manually curated, and spike time vectors were used to decode location - data contained in the local field potential (LFP) was discarded. Notably our CNN approach was consistently more accurate than the Bayesian decoder, exceeding its performance in all animals (Bayesian mean error 23.38cm ± 4.35cm; network error 17.31cm ± 4.46 cm; Wilcoxon signed-rank test: T=18, p=0.0001). Similarly, to compare the CNN against standard machine learning tools we used the wavelet transformed data to train support vector machines (SVMs). Note that in this case the spatial structure of the input is inevitably lost as the input features are transformed to a one-dimensional representation. Both linear (53.6cm ± 14.77cm; Figure 1E) and non-linear SVMs ((61.2cm ± 15.67cm) performed worse than the CNN.

The relative advantage over the Bayesian decoder increased further when the number of channels used for decoding was downsampled to simulate smaller recordings (linear regression Wald-test (n=31), s=−0.65, p=1.83e-10; Figure 1G). Notably, the model achieved a similar decoding performance with twenty tetrodes (80 channels, 23.45cm ± 3.15cm) as the Bayesian decoder reached with the full data set (128 channels, 23.25cm ± 2.79cm, Figure 1G). The high accuracy and efficiency of the model suggest that the CNN utilizes additional information contained in the LFP as well as from sub-threshold spikes and those that were not successfully clustered. Note that while the Bayesian decoder explicitly incorporates information about the animals’ positions at previous timesteps and probability with which each spatial location is visited, our model is effectively feed-forward - being presented with 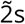 windows of data.

To better understand which elements of the raw neural data the network used, we retrained our model using datasets limited to just the LFP (<250Hz) and just the spiking data (>250Hz). In both cases, the network accurately decoded location (spikes-only (CNN-Spikes) mean error 17.23cm ± 4.69cm; LFP-only (CNN-LFP) mean error 24.24cm ± 6.00cm; Figure 1E), indicating that this framework is able to extract information from varied electrophysiological sources. Consistent with the higher information content of action potentials, the spikes-only network was considerably more accurate than the LFP-only network (Wilcoxon signed-rank test, two-sided (n=25): T=0, p=1.22e-05), although the LFP-only network was still comparable with the spike-based Bayesian decoder (Bayesian 23.38cm ± 4.35cm; LFP 24.24cm ± 6.00cm; Wilcoxon signed-rank test, two-sided (n=25): T=136, p=0.475). Note that previous studies have shown that demodulated theta is informative about the position of an animal in its environment (Agarwal et al., 2014). However, in those experiments theta oscillations were converted into a complex-valued signal, which carried both the magnitude and phase of theta here we only used the magnitude for decoding of position.

### Simultaneous decoding of multiple factors

The hippocampal representation of self-location is arguably one of the most readily identifiable neural codes - at any instance a small number of sparsely active neurons are highly informative. To provide a more stringent test of the network’s ability to detect and decode behavioural variables from unprocessed neural signals, we retrained with the same data but simultaneously decoded position, speed, and head-direction within a single model. CA1 recordings are known to incorporate information about these additional factors but their representation is less pronounced than that for self-location. Thus, the spatial activity of place cells is known to be weakly modulated by head direction (Jercog et al., 2019; Yoganarasimha et al., 2006), while place cell firing rates and both the frequency and amplitude of theta, a 7-10Hz LFP oscillation, are modulated by running speed (McFarland et al., 1975; Jeewajee et al., 2008). In this more complex scenario the architecture and hyper-parameters remained the same with just the final fully connected layer of the network being replicated, one layer for each variable, with the provision of appropriate loss functions - cyclical mean absolute error for head direction and mean absolute error for speed (see Methods). All three variables were decoded simultaneously and accurately (Position, 17.78cm ± 4.96cm; Head Direction, 0.80rad ± 0.18rad; Speed 4.94cm/s ± 1.00cm/s; Figure 2A & **Video 1**), with no meaningful decrement in performance relative to the simpler network decoding only position (position-only model 17.31cm ± 4.46; combined model 17.78cm ± 4.96cm; Wilcoxon signed-rank test two-sided (n=25): T=116, p=0.2108). Indeed, comparison of the *R*^2^-score metric from the fully trained network - a measure which represents the portion of variance explained and is independent of the loss function - indicated that mean decoding performance was above chance for all three behaviours (*R*^2^-score Position 0.86 ± 0.08, Head Direction 0.60 ± 0.12, Speed 0.72 ± 0.14, Chance *R*^2^-score Position −0.14 ± 0.13, Head Direction 0.04 ± 0.11, Speed −0.16 ± 0.22) (Figure 2B). Thus, the network was able to effectively access multiplexed information embedded in minimally processed neural data.

**Figure 2:**
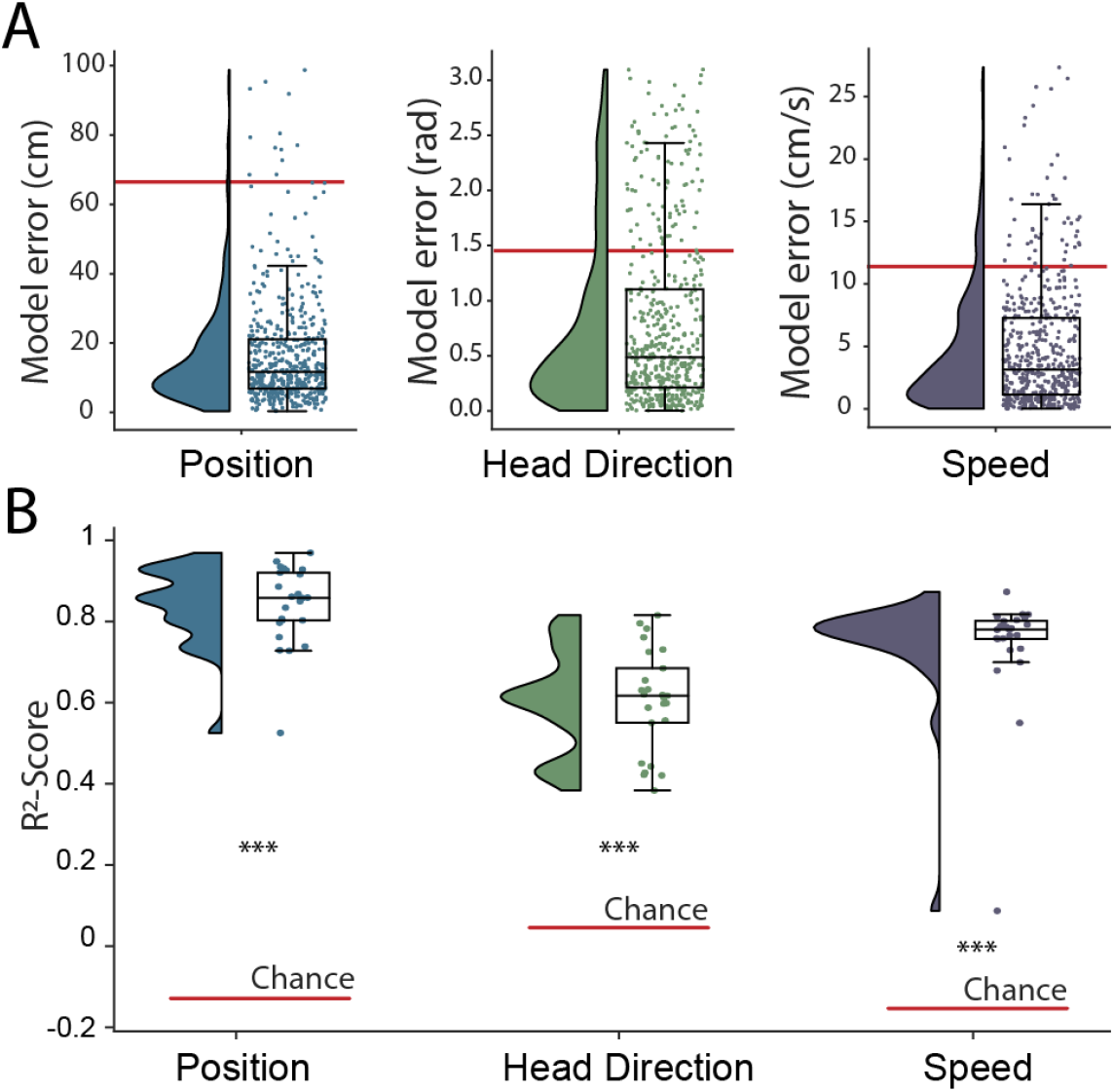
Simultaneous decoding of multiple variables from hippocampal data. A) Position, head direction, and running speed were accurately decoded in concert by a single network. Data from all five animals, each point indicates an error for a single sample. The red dashed line indicates the chance level obtained by shuffling the input relative to the output while fully retraining the model. B) *R*^2^-scores, a loss-invariant measure of model performance - ranging from 1 (perfect decoding) to negative infinity - allowing performance to be compared between dissimilar variables. Data as in (A), each point corresponds to one of five cross-validations within each of five rats.

### Interrogation of electrophysiological recordings

Although the network supports accurate decoding of self-location from electrophysiological data, this was not our main aim. Indeed, our primary goal for this framework was to provide a flexible tool capable of discovering and characterising sensory and behavioural variables represented in neural data - providing insight about the form and content of encoded information. To this end, in the fully trained network, we used a shuffling procedure to estimate the influence that each element of the 3D input (frequencies × channel × time) had on the accuracy of the decoded variables (see Methods). Since this approach does not require retraining it provides a rapid and computationally efficient means of assessing the contribution made by different channels, frequency bands, and time points.

Turning first to position decoding, we saw that the adjacent 469Hz and 663Hz frequency bands were by far the most influential - together accounting for more than 42% of the information about self-location derived from the electrophysiological data (Figure 3A). Since these recordings were made from CA1, we hypothesized that these frequencies corresponded to place cell action potentials. To confirm this hypothesis - and demonstrate that it was possible to objectively use this network-based approach to identify the neural basis of decoded signals - we applied the following approach (see Methods): First, we isolated the waveforms of place cells (n=629) and putative interneurons (n=91) in all animals, which were identified using a conventional approach (Pachitariu et al., 2016; Klausberger et al., 2003; Csicsvari et al., 1999). Second, for these two groups, we calculated the relative representation of the 26 frequency bands in their waveforms. We found that the highly informative 469Hz and 663Hz bands were the dominant components of place cell action potentials and that in general the power spectra of these spikes strongly resembled the frequency influence plot for position decoding (Spearman rank-order correlation, two-sided (n=26) ρ=0.84, p=7.63e-08; Figure 3B). In contrast, putative interneurons - which typically have a shorter after-hyperpolarisation than place cells (English et al., 2017) - were characterised by higher frequency components (Figure 3B, Mann-Whitney rank test interneuron (n=91) vs. place cell (n=629), U=1009.5, p=2.47e-13), with the highest power at 5304Hz and 3750Hz, bands that were considerably less informative about self-location (Figure 3A).

**Figure 3:**
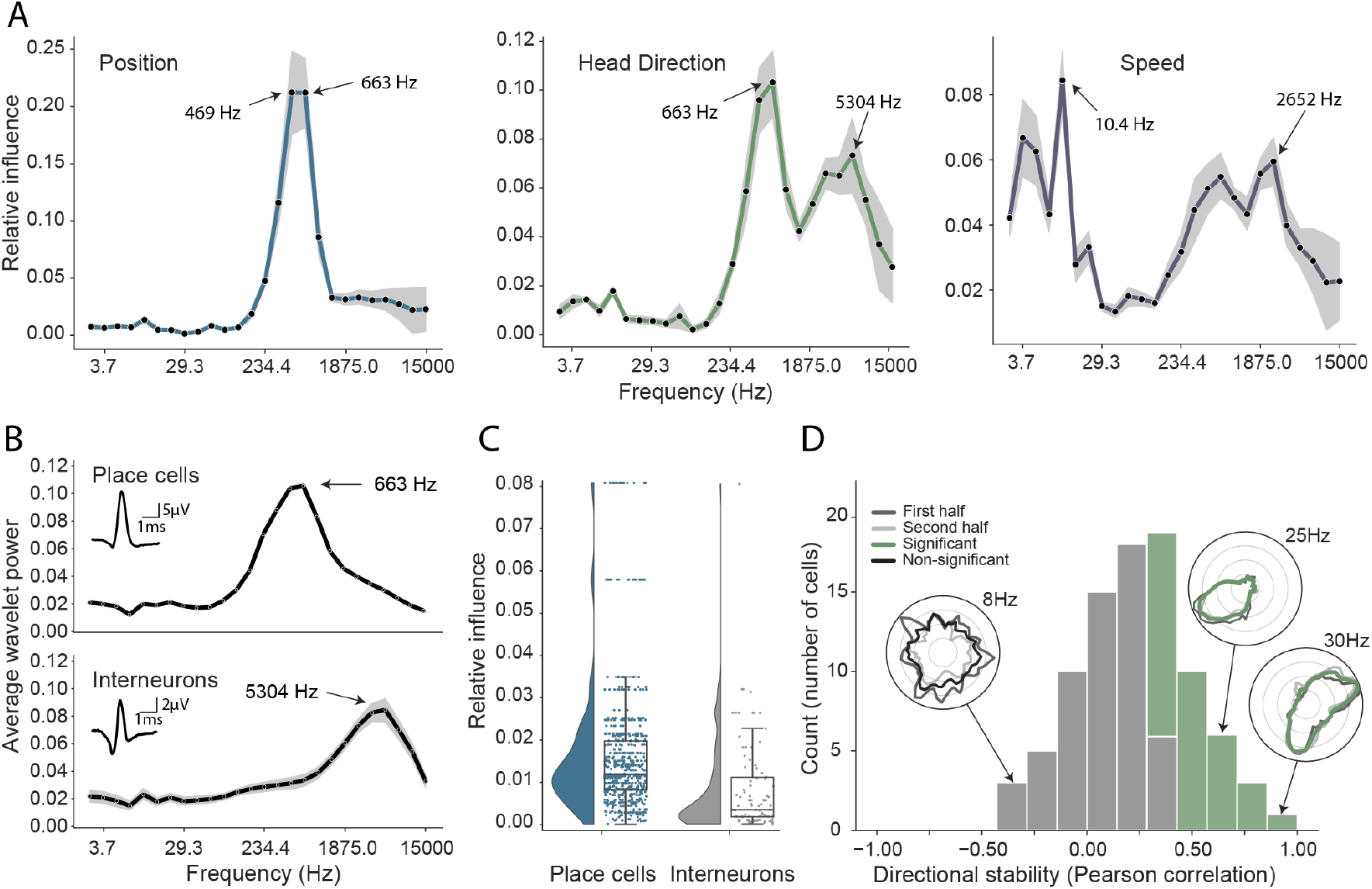
Analysis of trained network identifies informative elements of the neural code. (A) A shuffling procedure was used to determine the relative influence of different frequency bands in the network input. Left, the 469Hz and 663Hz components - corresponding to place cell action potentials - were highly informative about animals’ positions. Middle, both place cells and putative interneurons (5304Hz) carried information about head direction. Right, several frequency bands were informative about running speed, including those associated with the LFP (10.4Hz) and action potentials. Data from all animals. (B) Wavelet coefficients of place cell (top) and interneuron (bottom) waveforms are distinct and correspond to frequencies identified in A. Inset, average waveforms. Data from all animals. (C) Frequency bands associated with place cells (469 & 663Hz) were more informative about position than those associated with putative interneurons (5304 Hz) - their elimination produced a larger decrement in decoding performance (p<0.001). Data from all animals. (D) A subset of putative CA1 interneurons encodes head direction. 33/91 interneurons from five animals exhibited pronounced directional modulation that was stable throughout the recording (green). Depth of modulation quantified using Kullback-Leibler divergence vs. uniform circle. Stability assessed with the Pearson correlation between polar ratemaps from the first and second half of each trial (dark grey and light grey). Cells with p<0.01 for both measures were considered to be reliably modulated by head direction. Inset, example polar ratemaps. Data from all animals.

Since the frequencies associated with place cell waveforms were the most informative, this indicated that the network had correctly identified place cells as the primary source of spatial information in these recordings. To corroborate this, we used the same data and for each channel eliminated power in the 469Hz and 663Hz frequency bands at time points corresponding to either place cell or interneuron action potentials. As expected, position decoding was most strongly affected by removal of the place cell time points (Mann-Whitney-U-Test (n=629 place cells, n=91 interneurons): U=1497, p=2.86e-08; Figure 3C). Using the same shuffling method we also analysed how informative each channel was about self-location (Figure S3). In particular, we found that the number of place cells identified on a tetrode from the spike sorted data was highly correlated with the tetrode’s spatial influence (Spearman rank-order correlation (n=128) ρ=0.71, p=5.11e-06) and that the overall distribution of both number of place cells and spatial influence followed a log-normal distribution (Shapiro-Wilk test on log-transformed data, number of place cells, W=0.79, p=3.59e-05; tetrode influence W=0.59, p=3.04e-08; Figure S3B). In sum, this analysis correctly identified that the firing rates of both place cells and putative interneurons are informative about an animal’s location, place cells more so than interneurons (Wilent and Nitz, 2007). The analysis also highlighted the spatial activity of place cells, pointing to the stable place fields as a key source of spatial information.

A potential concern is that our approach might not identify multiple frequency bands if the information they contain is mutually redundant. The previous example, in which place cells and putative interneurons were both found to be informative about self-location, demonstrates this is not entirely the case. However, to further exclude this possibility we compared the influential frequencies identified from our complete model with models trained on just a single frequency band at a time. specifically, twenty-six models were trained, one for each frequency - the performance of each of these models being taken as an indication of the information present in that band. As expected we found both methods identified similar frequencies as indicated by a high correlation between our influence measure and the performance of single frequency band models (Position, Spearman rank-order correlation (n=26) ρ=0.88, p<0.001; Head Direction, ρ=0.82, p<0.001; Speed, ρ=0.47, p=0.02, Figure S4). Note that, although each model was individually faster to train than the complete model, the time to train all 26 was considerably longer than the single model applied simultaneously to all frequencies (51.2 hours vs. 8.6 hours, 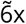 faster). Thus our combined approach provides a fair and efficient means to determine the informative elements of wide-band neural data. More importantly, analyses of the full network enables multiple frequency bands to be considered in-concert, providing a means to identify interactions (e.g.Figure S5) that are not accessible to standard single-frequency methods.

### CA1 interneurons are modulated by head direction

Next, having validated our approach for spatial decoding, we examined the basis upon which the network was able to decode head direction. Although place cells primarily provide an allocentric spatial code, their infield firing rate is known to be modulated by heading direction (Muller et al., 1994; Rubin et al., 2014). Consistent with the presence of this directional code, we again saw that the most influential frequencies for head direction decoding were those associated with place cells (469Hz and 663Hz; Figure 3A). However, the distribution also incorporated a secondary peak corresponding to the frequencies typical for interneuron waveforms (Spearman rank-order correlation (n=26) ρ=0.76, p=5.71e-06, Fig 3AB). Presubicular interneurons have been shown to be modulated by both head direction and angular velocity (Preston-Ferrer et al., 2016) but to the best of our knowledge no similar responses have been noted in CA1. To establish if putative interneurons conveyed information about head direction we again used an ‘elimination’ analysis on data from all five animals - the two frequency bands most strongly associated with interneurons (3750Hz and 5304Hz) were scrambled at time points when interneuron spikes were present. Consistent with the influence plots, we found that selectively eliminating putative interneurons degraded the accuracy with which head direction was decoded (relative influence: 0.089 ± 0.043, two-sided t-test (n=91) t=4.16, p=0.014). As a final step, to verify this novel observation we reverted to a standard approach. specifically, we calculated the directional ratemap for each interneuron using only periods when the animal was in motion (>10cm/s), determined the Kullback-Leibler divergence vs. a uniform circle (Doeller et al., 2010), and applied a shuffling procedure to determine significance - as a whole the population exhibited reliable but weak modulation of interneuronal firing rate by head direction (Kullback-Leibler Divergence (n=91): 0.0067 ± 0.009) with 58.2% (53/91) of cells being individually significant (p<0.01). Behaviours that are inhomogeneously distributed or confounded can result in spurious neural correlates (Muller et al., 1994). To control for this possibility we repeated the analysis using only data from the centre of the environment (>25cm from the long sides of the enclosure and >20cm from the short sides). Additionally, to verify stability, we controlled that ratemaps generated from the first and second half of the trial were correlated (Pearson correlation, p<0.01). Under this more rigorous analysis, we confirmed that a sub-population (36.2%, 33/91) of putative hippocampal interneurons were modulated by head direction, a previously unrecognised spatial correlate (Figure 3D).

### Multiple electrophysiological features contribute to the decoding of speed

The frequency influence plots for running speed also showed several local peaks (Figure 3A), that in all cases corresponded to established neural correlates. In rodents, theta frequency and power are well known to co-vary almost linearly with running speed (McFarland et al., 1975; Jeewajee et al., 2008), accordingly analysis of the network identified the 10.4Hz frequency band as the most influential. Similarly, the firing rate of place cells increases with speed, an effect captured by the peak at 663Hz. Interestingly a clear peak is also evident at 2652Hz, indicating that interneuron firing rates are also informative - originating either from CA1 speed cells (Góis and Tort, 2018) or from theta-locked interneurons (Huh et al., 2016). Finally, a 4th peak was evident at 3.66Hz and 5.17Hz, a range that corresponds to type 2 (’atropine sensitive’) theta which is present during immobility (Kramis et al., 1975; Sainsbury et al., 1987). To corroborate this conclusion, we calculated the correlation between power in each frequency band and running speed (Figure S6A), confirming that the latter band showed the expected negative correlation - higher power at low speeds - while the other three peaks were positively correlated.

### Generalization across brain regions and recording techniques

As a final step, we sought to determine how well our approach generalised to other recording techniques and brain areas. Addressing the latter point first, we trained the network using electrophysiological recordings (64 channels) from the primary auditory cortex of a freely-moving mouse while pure tone auditory stimuli (4 to 64kHz, duration 200 ms) were played from a speaker (Figure 4A). As above, the raw electrophysiological data was transformed to the frequency domain using Morlet wavelets and this wide-band frequency representation was used as input. The model architecture and hyperparameters were kept the same, reducing only the number of down-sampling steps because of the smaller input size(64 channels vs. 128 channels for CA1 recordings - each down-sampling layer halves the number of units in previous layer). The auditory stimuli - training target - was modelled as a continuous variable with ‘−1’ indicating no tone present and the log-transformed frequency of the sound at all other time points. As expected, this model architecture was also able to accurately decode tone stimuli from auditory cortex (*R*^2^-scores of 0.734 ± 0.080, chance model: −0.432 ± 0.682, Figure 4B,C). Informative frequencies were concentrated around 663Hz and 165Hz, indicating that information content about tone stimuli comes mostly from pyramidal cell activity.

**Figure 4:**
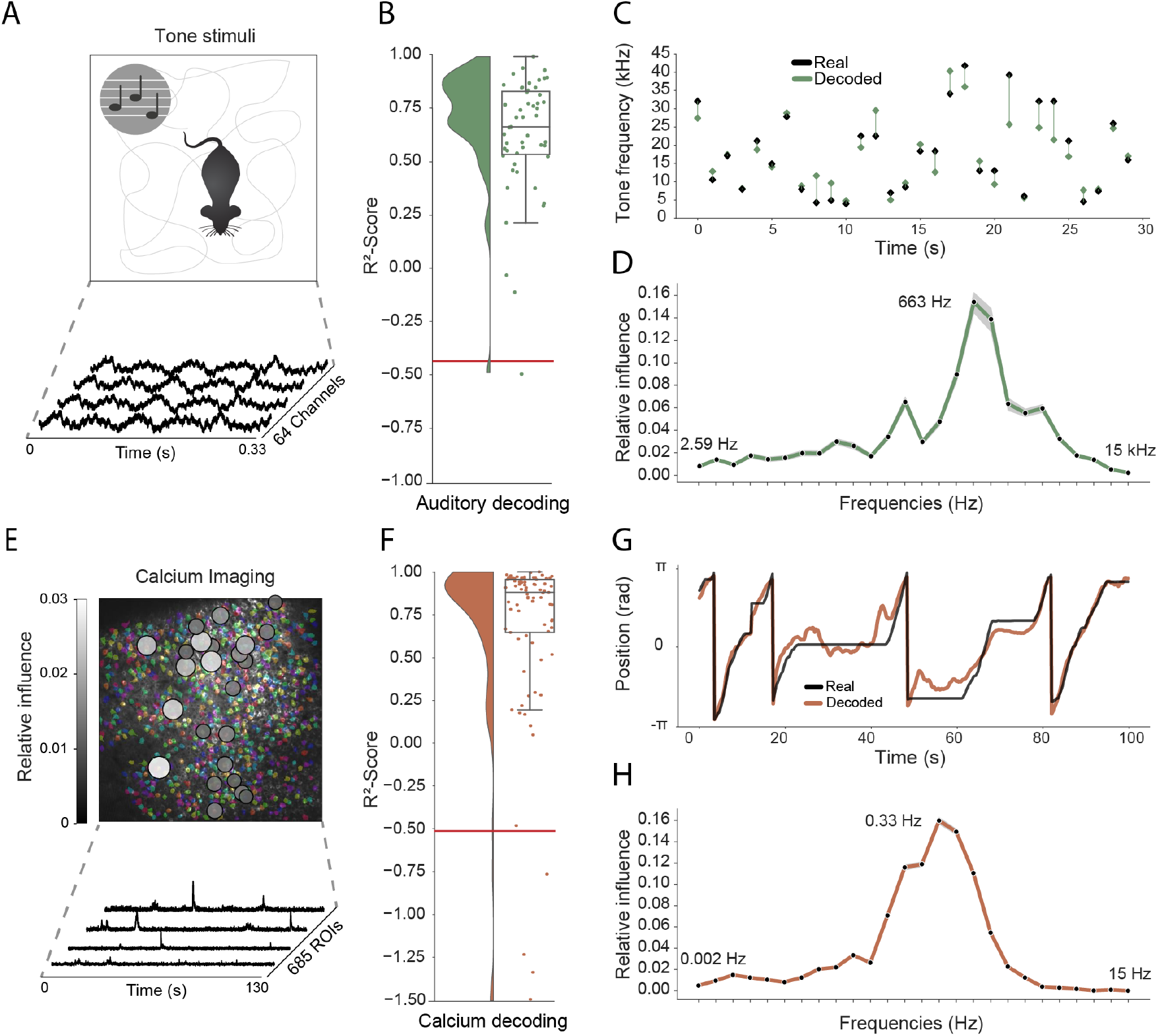
Model generalizes across recording techniques and brain regions. (A) Overview of auditory recording. We recorded electrophysiological signals while the mouse is freely moving inside a small enclosure and is presented with pure tone stimuli ranging from 4kHz to 64kHz. (B) *R*^2^-score for decoding of frequency tone from auditory cortex (0.73 ± 0.08). Each dot describes the *R*^2^-score for a 5s sample of the experiment. Chance level is indicated by the red line. (C) An example section for decoding of auditory tone frequencies from auditory electrophysiological recordings, real tone colored in black, decoded tone in green, the line between real and decoded indicates magnitude of error. (D) influence plots for decoding of auditory tone stimuli, same method as used for CA1 recordings. (E) Calcium recordings from a mouse running on a linear track in VR. We record from 685 cells and use Suite2p to preprocess the raw images and extract calcium traces which we feed through the model to decode linear position. Overlay shows relative influence for decoding of position calculated for each putative cell. (F) *R*^2^-score for decoding of linear position from two-photon CA1 recordings (0.90 ± 0.03). Each dot describes the *R*^2^-score for a 5s sample of the experiment. Chance level is indicated by the red line. (G) Example trajectory through the virtual linear track (linearized to [−*π, π*] with real position (black) and decoded position (orange)). (H) influence plots for decoding of position from two-photon calcium imaging. Note that the range of frequencies is between 0Hz and 15Hz as the sampling rate of the calcium traces is 30Hz.

Having shown that the model generalises across different brain areas we wanted to further investigate if it generalizes across different recording techniques. Therefore, in the third set of experiments we acquired two-photon calcium fluorescence data from mouse CA1 while the head-fixed animal explored a 230cm virtual track. Raw data was preprocessed to generate denoised activity traces for putative cells (n=685 regions of interest), these were then decomposed to a frequency representation using the same wavelet approach as before - only frequency bands between 0Hz and 15Hz being used because of the lower data rate (30Hz) (see methods, Figure 5E). As before, wavelet coefficients were provided to the network as input and the only change was an increase in the number of down-sampling steps to account for the large number of ROIs (685 ROIs vs 128 channels for CA1 recordings). The network was able to accurately decode the animal’s position on the track (mean error: 15.87 ± 16.33 cm, *R*^2^-scores of 0.90 ± 0.03 vs. chance model −0.05 ± 0.127, Figure 4F,G). Using the same shuffling technique as before, we generated influence plots indicating the relative information provided by putative cells (Fig 4E) and frequencies (Fig 4H). In the frequency domain, the most informative bands were 0.33Hz and 0.46Hz, unsurprisingly mirroring the 1s to 2s decay time of GCaMP6s (Chen et al., 2013). Interestingly the relative information content of individual cells was highly heterogeneous, a small subset (18.2%) of cells accounted for half (50%) of the influence - these units being distributed across the field of view with no discernible pattern (Figure 4E).

## Discussion

The neural code provides a complex, non-linear representation of stimuli, behaviours, and cognitive states. Reading this code is one of the primary goals of neuroscience - promising to provide insights into the computations performed by neural circuits. However, decoding is a non-trivial problem, requiring strong prior knowledge about the variables encoded and, crucially, the form in which they are represented. Not only is this information often incomplete or absent but a full characterisation of the neural code is precisely the question we seek to solve. Addressing these limitations, we investigated the potential of a deep-learning framework to decode behaviours and stimuli from wide-band, minimally processed neural activity. To this end, we designed a model architecture using simple 2D convolutions with shared weights, omitting recurrent layers (Bai et al., 2018). These intentional design choices resulted in a fast, data efficient architecture that could be easily interpreted to discover which elements of the neural code provided information about specific variables - a decrease in network performance was accepted as a trade-off. We showed that this approach generalised well across brain regions and recording techniques capturing both spatial and temporal information in the signal, the only changes necessary to the network being adjustments to handle the number of channels in the input matrix.

In the first instance we validated our model using the well characterised spatial representations of rodent CA1 place cells. Decoding performance amply exceeded a Bayesian framework, as well as a standard machine learning approach that proved ineffective on the non-linear representation of self-location. Importantly, simple analyses of the trained network correctly indicated that place cell action potentials were the most informative spatial signal - confirming that this tool can deliver insights into the nature of the neural code. In a further set of experiments we showed that the network was able to concurrently identify multiple representations of head direction and running speed, including several that were only recently reported and one - interneuron encoding of head direction - that was previously unreported. Importantly, this framework can also identify interactions between frequency components, an analysis that is intractable to conventional methods which consider features independently. Finally, we demonstrated the flexibility of this approach, applying the same network and hyper-parameters, with adjustments made only to the input and output layers, to two-photon calcium data and extracellular recordings from auditory cortex.

In sum, we believe deep-learning based frameworks such as this constitute a valuable tool for experimental neuroscientists, being able to provide a general overview as to whether a variable is encoded in time-series data and also providing detailed information about the nature of that encoding - when, where, and in what frequency bands it is present. That is not to say that this approach is a complete substitute for conventional analyses - it merely constrains the search space for variables that might be present and their plausible format. Indeed, we imagine this network might be best used as a first pass analysis, followed by conventional approaches to determine explicitly if a variable is present - much as we did for the interneuron representation of head direction. While we tested the network with optical and electrophysiological data it is highly likely that it will perform well with neural data acquired in most experimental settings, including fMRI, EEG, and MEG.

## Methods

### Tetrode recordings from CA1

Five male Lister Hooded rats were used for this study. All procedures were approved by the UK Home Office, subject to the restrictions and provisions contained in the Animals Scientific Procedures Act of 1986. All rats (333-386 g/13-17 weeks old at implantation) were implanted with two single-screw microdrives (Axona Ltd.) targeted to the right and left CA1 (ML: 2.5 mm, AP: 3.8 mm posterior to bregma, DV: 1.6 mm from dura). Each microdrive was assembled with two 32 channel Omnetics connectors (A79026-001) and 16 eight tetrodes of twisted wires (either 17 *μ*m H HL coated platinum iridium, 90% and 10% respectively, or 12.7 *μ*m HM-L coated Stablohm 650; California Fine Wire), platinum plated to reduce impedance to below 150 kΩ at 1 kHz (NanoZ). After surgery, rats were housed individually on a 12 hr light/dark cycle and after one week of recovery rats were maintained at 90% of free-feeding weight with ad libitum access to water.

Screening was performed from one week after surgery. Electrophysiological data was acquired using Open Ephys recording system (Siegle et al., 2017) and a 64-channel amplifier board per drive (Intan RHD2164). Positional tracking performed using a Raspberry Pi with Camera Module V2 (synchronised to Open Ephys system) and custom software, that localised two different brightness infra-red LEDs attached to amplifier boards on camera images acquired at 30 Hz. During successive recording sessions in a separate screening environment 1.4 × 1.4 m the tetrodes were gradually advanced in 62.5 *μ*m steps until place cells were identified. During the screening session, the animals were often being trained in a spatial navigation task for projects outside the scope of this study.

The experiments were run during the animals’ dark period of the L/D cycle. The recording sessions used in this study were around 40 min long, depending on the spatial sampling of the animal, in a rectangular environment of 1.75 × 1.25 m, on the second, third or fourth exposure, varying between animals. The environment floor was black vinyl flooring, it was constructed of 60 cm high boundaries (MDF) colored matt black, surrounded by black curtains on the sides and above. There was one large cue card raised above the boundary and two smaller cue cards distributed on the side of the boundary. Foraging was encouraged with 20 mg chocolate-flavoured pellets (LBS Biotechnology) dropped into the environment by custom automated devices. The recordings used in this study were part of a longer session that involved foraging in multiple other different size open field environments.

Rats were anaesthetised with isoflurane and given intraperitoneal injection of Euthanal (sodium pentobarbital) overdose (0.5 ml / 100 g) after which they were transcardially perfused with saline, followed by a 10% Formalin solution. Brains were removed and stored in 10% Formalin and 30% Sucrose solution for 3-4 days prior to sectioning. Subsequently, 50 *μ*m frozen coronal sections were cut using a cryostat, mounted on gelatine coated or positively charged glass slides, stained with cresyl violet and cleared with clearing agent (Histo-Clear II), before covering with DPX and coverslips. Sections were then inspected using Olympus microscope and tetrode tracks reaching into CA1 pyramidal cell layer were verified.

Putative interneurons were classified based on waveform shape, minimum firing rate across multiple environments and lack of spatial stability. specifically, classi1ed interneurons had waveform half-width less than 0.15 ms, maximum ratio of amplitude to trough of 0.4, minimum firing rate of 4 Hz and maximal 0.75 spatial correlation of ratemaps from first and last half of the recording in any environment (Klausberger et al., 2003; Csicsvari et al., 1999). Note that we used the spatial stability in order to differentiate interneurons from place cells or grid cells, with no influence on the directional stability of the head direction cell analysis.

### Calcium recordings from CA1

All procedures were conducted in accordance to UK Home Office regulations

One GCaMP6f mouse (C57BL/6J-Tg(Thy1-GCaMP6f)GP5.17Dkim/J, Jacksons) was implanted with an imaging cannula (a 3mm diameter × 1.5mm height stainless-steel cannula with a glass coverslip at the base) over CA1 (stereotaxic coordinates: AP=−2.0, ML=−2.0 from bregma). A 3mm craniotomy was drilled at these coordinates. The cortex was removed via aspiration to reveal the external capsule of the hippocampus. The cannula was inserted into the craniotomy and secured to the skull with dental cement. A metal head-plate was glued to the skull and secured with dental cement. The animal was left to recover for at least one week after surgery before diet restriction and habituation to head-fixation commenced.

Following a period of handling and habituation, the mouse was head-fixed above a styrofoam wheel and trained to run for reward through virtual reality environments, presented on 3 LCD screens that surrounded the animal. ViRMEn software (Aronov and Tank, 2014) was used to design and present the animal with virtual reality linear tracks. Movement of the animal on the wheel was recorded with a rotatory encoder and lead to corresponding translation through the virtual track. During the experimental phase of the training, the animal was trained to run down a 230cm linear track and was required to lick at a reward port at a fixed, unmarked goal location within the environment in order to trigger release of a drop of condensed milk. Licks were detected by an optical lick detector, with an IR LED and sensor positioned on either side of the animal’s mouth. When the animal reached the end of the linear track, a black screen appeared for 2 seconds and the animal was presented with the beginning of the linear track, starting a new trial.

Imaging was conducted using a two-photon microscope (resonant scanning vivoscope, Scientifica) using 16x/0.8-NA water-immersion objective (Nikon). GCaMP was excited using a Ti:sapphire laser (Mai Tai HP, Spectra-Physics), operated with an excitation wavelength of 940nm. ScanImage software was used for data collection / to interface with the microscope hardware. Frames were acquired at a rate of 30Hz.

The Suite2p toolbox (Pachitariu et al., 2017) was used to motion correct the raw imaging frames and extract regions of interest, putative cells.

### Tetrode recordings from auditory cortex

Sound-evoked neuronal responses were obtained via chronically-implanted electrodes in the right hemisphere auditory cortex of one 17-week-old male mouse (M. musculus, C57Bl/6, Charles River). All experimental procedures were carried out in accordance with the institutional animal welfare guidelines and a UK Home Office Project License approved under the United Kingdom Animals (Scientific Procedures) Act of 1986.

During recordings, the animal was allowed to freely move within a 11×21 cm cardboard enclosure, with one wall consisting of an acoustically-transparent mesh panel to allow unobstructed sound stimulation. Acoustic stimuli were delivered via two free-field electrostatic speakers (Tucker-Davis Technologies, FL, USA) placed at ear level, 7 cm from the edge of the enclosure. Recordings were performed inside a double-walled soundproof booth (IAC Acoustics), whose interior was covered by 4-cm thick acoustic absorption foam (E-foam, UK). Pure tones were generated using MATLAB (Matlab version R2015a; MathWorks, Natwick, MA, USA), and played via a digital signal processor (RX6, Tucker Davis Technologies, FL, USA). The frequency response of the loudspeaker was ±10 dB across the frequency range used for stimulation. Pure tones of a duration of 200ms (including 5 ms linear rise and fall times) of variable frequencies (4-64 kHz in 0.1 octave increments) were used for stimulation. The tones were presented at 65dB SPL at the edge of the testing box. The 41 frequencies were presented pseudo-randomly, separated by a randomly varying inter-stimulus interval ranging from 500 to 510ms, for a total of 20 repetitions.

Extracellular electrophysiological recordings were obtained using a custom chronically-implanted 64-channel hyperdrive with two 32-channel Omnetics connectors (A79026-001) and 16 individually movable tetrodes (FlexDrive, (Voigts et al., 2013)). Tetrodes were made from 12.7 *μ*m tungsten wire (99.95%, HFV insulation, California Fine Wire, USA) gold-plated to reduce impedance to 200kΩ at 1kHz (NanoZ, Multichannel Systems). Neuronal signals were collected and amplified using two 32-channel amplifier boards (Intan RHD 2132 headstages) and an Open Ephys recording system (Siegle et al., 2017) at 30kHz.

### Data Preprocessing

Raw electrophysiological traces as well as calcium traces were transformed to a frequency representation using discrete-wavelet transformation (DWT). We decided to use wavelet transformation instead of windowed Fourier transform (WFT) as we expected a wide range of dominant frequencies in our signal for which the wavelet transformation is more appropriate (Torrence and Compo, 1998). For the wavelet transformation, we used the morlet wavelet:

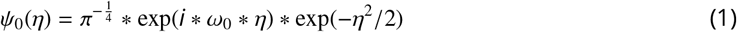

with a non-dimensional frequency constant *w*_0_ = 6. We noticed that downscaling the wavelets improved our model performance, prompting us to use an additional preprocessing step which effectively decreased the sampling rate of the wavelets to 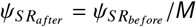 by a factor of *M*. This can also be seen as an additional convolutional layer with a kernel size of *M*, a stride of *M* and weights fixed to 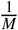. We performed a hyperparameter search for *M* with a simplified model and found the best performing model with *M* = 1000, thus effectively decreasing our sampling rate from 30000 to 30 (Figure S7).

As additional preprocessing steps we applied channel and frequency wise normalization using a median absolute deviation (MAD) approach. We calculated the median and the corresponding median absolute deviation for each frequency and channel on the training set and normalized our inputs as follows:

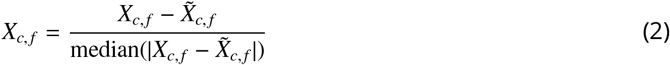

where 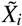 is the median of *X_i_*. This approach turned out to be more robust against outliers in the signal than simple mean normalization. Additional min-max scaling did not further improve performance.

### Bayesian decoder

As a baseline model, we used a Bayesian decoder which was trained on manually sorted and clustered spikes. Given a time window *T* and number of spikes *K* = (*k*_1_, …*k_n_*) fired by *N* place cells, we can calculate the probability *p*(*K*|*x*), estimating the number of spikes *K* at location *x*:

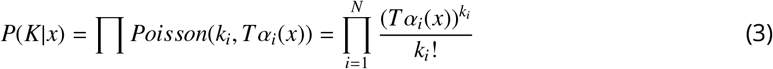

where *α_i_*(*x*) is the firing rate of cell *i* at position *x*. From this we can calculate the probability of the animals location given the observed spikes:

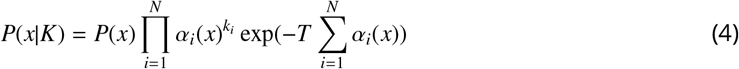

where *P*(*x*) is the historic position of the animal which we use to constrain *p*(*x*|*K*) to provide a fair comparison to the convolutional decoder. The final estimate of position is based on the peak of *p*(*x*|*K*):

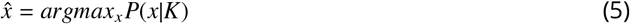

We used the same cross-validation splits as for the convolutional model and calculated the Euclidean distance between the real and decoded position. We performed grid search on one representative rat to 1nd the optimal parameters regarding bin size and bin length for the Bayesian decoder. The optimized Bayesian decoder uses a Gaussian smoothing kernel with sigma = 1.5, a bin size of 2cm for binning the ratemaps, and uses a bin length of 0.5s.

### Convolutional neural network

The model takes a three-dimensional wavelet transformed signal as input and uses convolutional layers connected to a regression head to decode continuous behaviour. We use a kernel size of 3 throughout the model and keep the number of filters constant at 64 for the first 8 layers while sharing the weights over the channel axis, while then doubling them for each following layer. For downsampling the input we use a stride of 2, intermixed between the time and frequency dimension (Table S1, Fig 1B). As regularization we apply gaussian noise 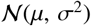 with *μ* = 0 and *σ* = 1 to each input sample.

We extensively investigated the use of a convolutional long-short-term-memory (LSTM) after the initial convolutions, where we used backpropagation through time on the time dimension. In a simplified model this led to a small decrease in decoding error for the model trained on position. We nevertheless decided to employ a model with only simple convolutions as one important aspect of this model is the simplicity of use for a neuroscientist. Moreover we experimented with using a wavenet (Oord et al., 2016) inspired model directly on the raw electrophysiological signal but noticed that the model using the wavelet transformed input outperformed the wavenet approach by a margin of around 20cm for the positional decoding. The wavenet inspired model was considerably slower to train and therefore a full hyperparameter search could not be performed.

Previous models contrasting recurrent vs. convolutional networks (Bai et al., 2018), find that convolutional layers outperform recurrent ones when trained directly on minimally processed data. The benchmarks typically used in classical sequence learning are one-dimensional, whereas we record two-dimensional raw input (time × channels) with a high sampling rate, complicating the amount of experimentation we could perform as the unprocessed data for a 2s time window exceeds the capacity of GPU memory (30000 × 128 time points per sample). In the related field of speech processing with sampling rates up to 48000Hz, the input is processed using log-mel feature banks which are computed with a 25ms window and a 10ms shift (Bahdanau et al., 2016; Chan et al., 2016; Prabhavalkar et al., 2017). We therefore opted for a similar approach by using downsampled wavelet transformed signals, resulting in a 33.3ms window given a downsampling size of z=1000. Note that with further downsampling there might be a risk of losing decoding precision, with some of the behaviours coming close to the downsampled rate (e.g. head direction can be up to 40deg/s) (Figure S7).

### Model training

The model takes as input a three-dimensional wavelet transformed signal corresponding to time, frequency and channels, with frequencies logarithmically scaled between 0Hz-15000Hz. An optimal temporal window of T=64 (corresponding to 2.13s) was established by hyperparameter search taking into account the tradeoff between speed of training and model error. For training the model across the full duration of the experiment we divided the experiment into 5 partitions and used cross-validation for testing the model on before unseen data partitions, i.e. we first used partitions 2 to 5 for training and 1 for testing, then 1,3,4 and 5 for training and 2 for testing and so on. The last partition uses 1 to 4 for training and 5 for testing. Importantly, the overlap introduced by using 2s long samples was accounted for by using gaps (2s) between the training partitions, making sure that training and test set are fully independent of each other. We then randomly sampled inputs and outputs from the training set. Each input had corresponding outputs for the position (X, Y in cm), head direction (in radians) and speed (in cm/s). We used Adam as our learning algorithm with a learning rate of 0.0007 and stopped training after we sampled 18000 samples, divided into 150 batches for 15 epochs, each batch consisting of 8 samples. During training we multiplied the learning rate by 0.2 if validation performance did not improve for 3 epochs. We performed random hyperparameter search for the following parameters: learning rate, dropout, number of units in the fully connected layer and number of input timesteps. For calculating the chance level we used a shuffling procedure in which the wavelet transformed electrophysiological signal is shifted relative to its corresponding position. After shuffling we trained the model with the same setting as the unshuffled model and for the same number of epochs. The training was performed on one GTX1060 using Keras with Tensorflow as backend.

### Model comparison

In order to compare the performance of the network against the Bayesian decoder we simulated both models in a setting with artificially reduced inputs. We used 1 to 32 tetrodes as input for both decoders, with tetrodes taken top to bottom in order of the given tetrode number. The input of run 1 was then comprised of tetrodes 1 to 32, while run 2 used tetrodes 1 to 31. The last run uses only the first tetrode as input to both models. We then retrained both models with the artificially reduced number of tetrodes making sure both models have the same cross-validation splits and report decoding errors as the average of each cross-validation split.

### Model evaluation

For adjusting the model weights during training we use different loss terms depending on the behaviour or stimuli which we decode.

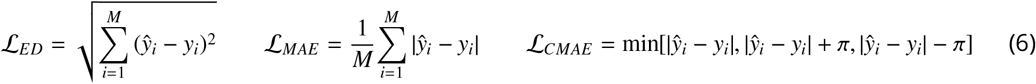

For decoding of position from tetrode CA1 recordings we try to minimize the Euclidean loss between predicted and ground truth position 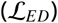. We use the mean squared error for the decoding of speed 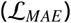 and the cyclical absolute error for decoding of head direction 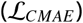. For all other behaviours or stimuli we use 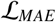 as the default optimizer.

We decided to use *R*^2^ scores to measure model performance across different behaviours, brain areas and recording techniques. We use the formulation of fraction of variance accounted for (FVAF) instead of the squared Pearson’s correlation coefficient. Both terms are based on the fraction of the residual sum of squares and the total sum of squares:

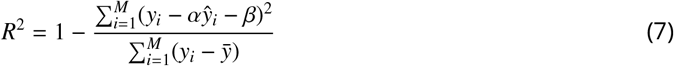

with *y_i_* the ground truth of sample *i*, 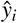 the predicted value and 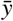 the mean value. Here, Pearson’s correlation coefficient tries to maximize *R*^2^ by adjusting α and β while FVAF uses α = 1 and β = 0 (Fagg et al., 2009). This provides a more conservative measure of performance as FVAF requires that prediction and ground truth fit without scaling the predicted values. FVAF in turn has no lower bound as the prediction can be arbitrarily worse with a given scaling constant (i.e. given a ground truth value of 10, a prediction of 1000 has a lower (worse) *R*^2^ score than a prediction of 100).

### influence maps

To investigate which frequencies, channels or timepoints were informative for the respective decoding we performed a bootstrapping procedure after training the models. For each sample in time we calculated the real decoding error *e_o_* for each behaviour by using the wavelets as input. We then shu[e the wavelets for a particular frequency and re-calculate the error. We then define the influence of a given frequency or channel as the relative change: 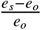 where *e_o_* is the original error and *e_s_* the shu[ed error. We repeat this for the channel and time dimension to get an estimate of how much influence each channel or timepoint has on the decoding of a given behaviour.

We also tried calculating sample gradients with respect to our inputs (Simonyan et al., 2013). For this we calculated the derivative *w* by back-propagation for each sample and with respect to the inputs. In contrast to class saliency maps, we obtain a gradient estimate indicating how much each part of the input strongly drives the regression output. We calculate saliency maps for each sample cross-validated over the entire experiment. For deriving influence maps from the raw gradients we calculate the variance across the time dimension and use this as an estimate of how much influence each frequency band or channel has on the decoding. This method however introduces a lot of high-frequency noise in the gradients, possibly coming from the strides in the convolutional layers used throughout the model (Olah et al., 2017).

## Author Contributions

CB, JK initial idea for project, MF conceived the network architecture and implemented all neural network models, CB implemented the Bayesian decoder, ST spike sorted the data and classified neurons, ST, CP, AL acquired data with help from DB, MF, ST, CB, MN, AB, JK, CFD contributed ideas to experiments and analysis, MF, CB wrote the paper with input from all authors.

MF and CFD are supported by the European Research Council (ERC-CoG GEOCOG 724836); CFD’s research is also funded by the Max Planck Society; the Kavli Foundation; the Centre of Excellence scheme of the Research Council of Norway – Centre for Neural Computation (223262/F50), The Egil and Pauline Braathen and Fred Kavli Centre for Cortical Microcircuits, and the National Infrastructure scheme of the Research Council of Norway – NORBRAIN (197467/F50); CB funded by Wellcome SRF (212281/Z/18/Z); CP funded by Wellcome Trust (110238/Z/15/Z).

## Data Availability

All datasets used in this study will be made public upon publication.

## Code Availability

The code is available at **https://github.com/CYHSM/DeepInsight**

## Supplementary Material

**Figure S1:**
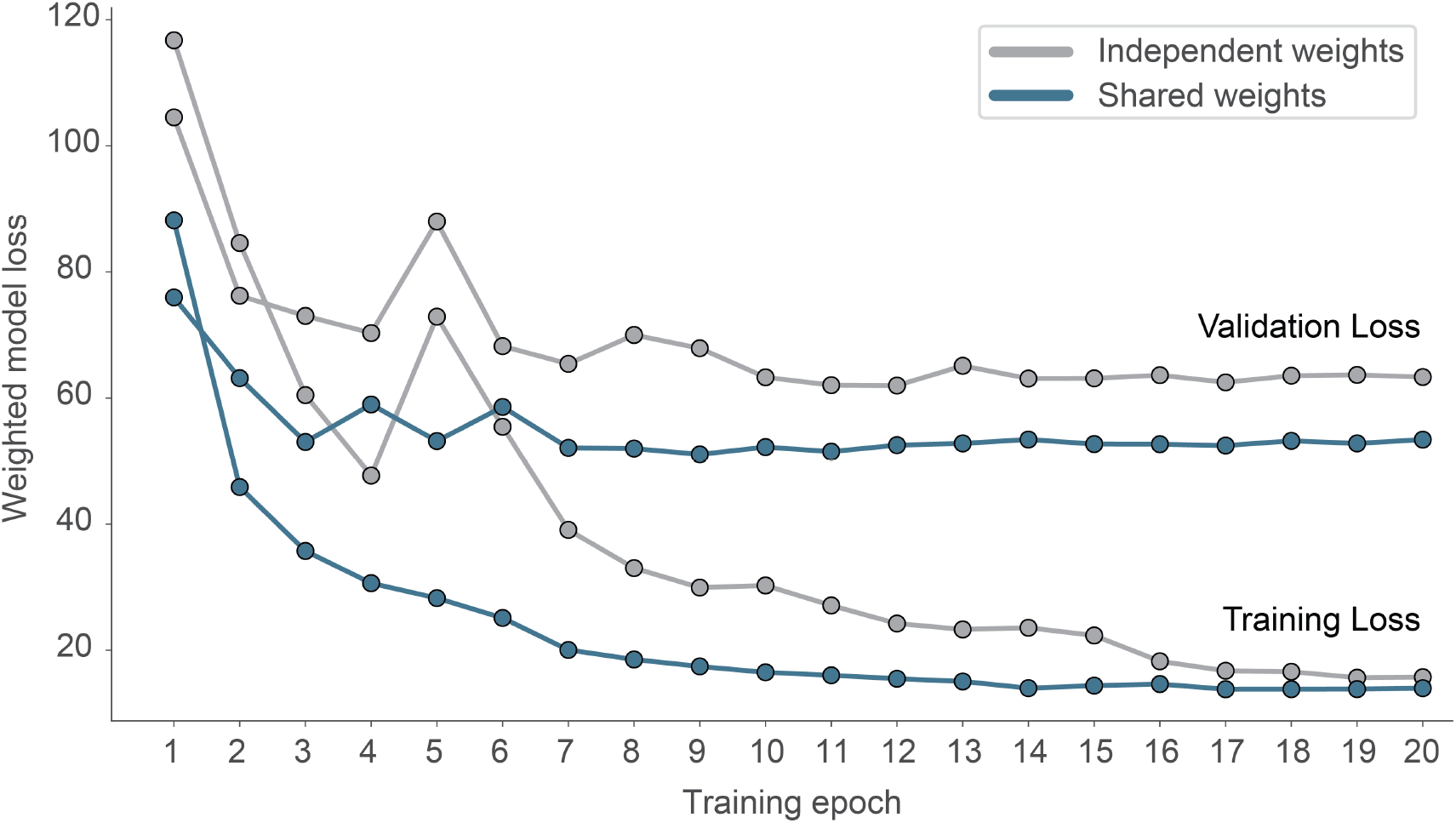
Effect of weight sharing on model performance. We evaluated two models on the same dataset, using either shared weights and 2D-convolutions (blue) or independent weights using 3D-convolutions (grey). The model using shared weights reaches a lower validation loss and generalizes better (smaller overfit).

**Figure S2:**
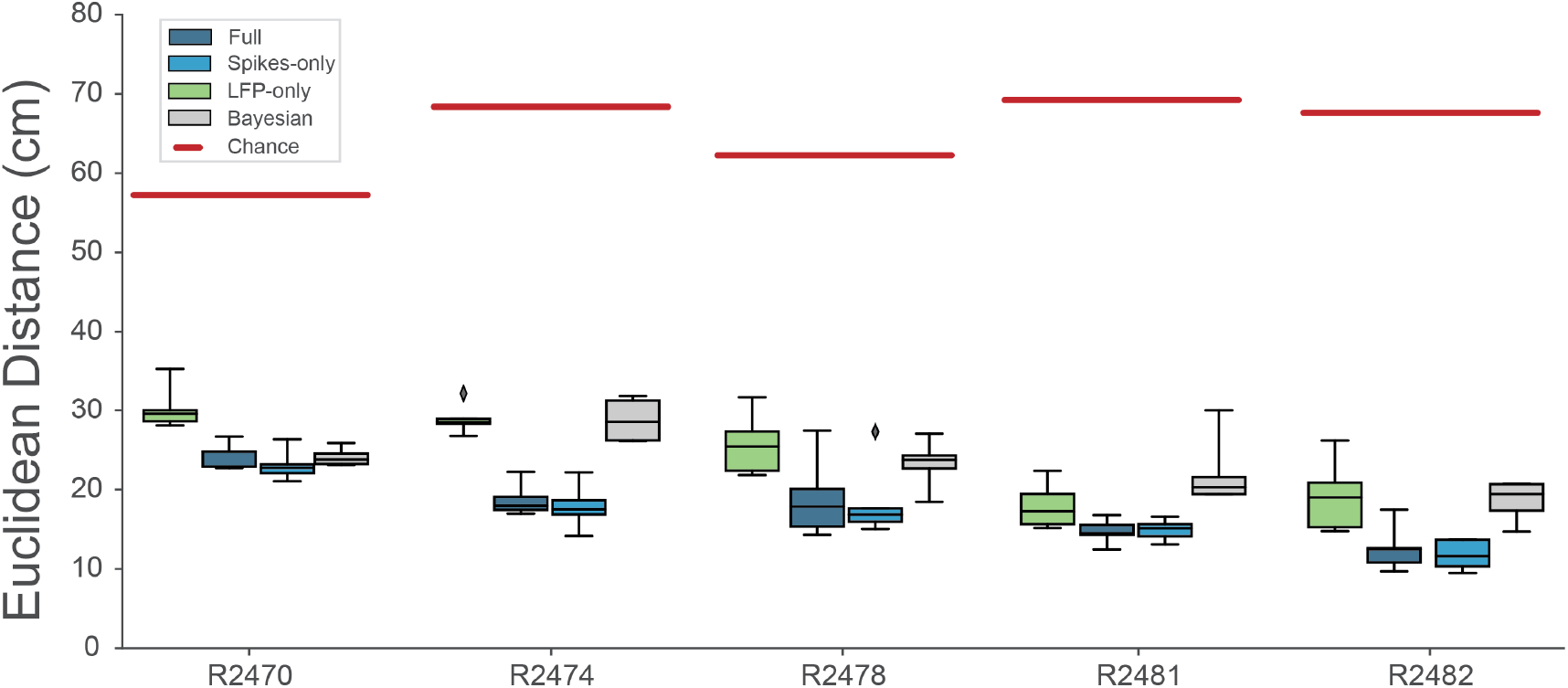
Decoding performance across different models. We calculated the Euclidean distance between the real behaviour and decoded behaviour across five rats and 4 different models. Full model has access to all frequency bands from 2-15000Hz, Spikes model has access to frequencies >250Hz, while LFP model uses frequencies <250Hz. Bayesian decoder was trained on spike sorted data. Chance level is indicated as red line.

**Figure S3:**
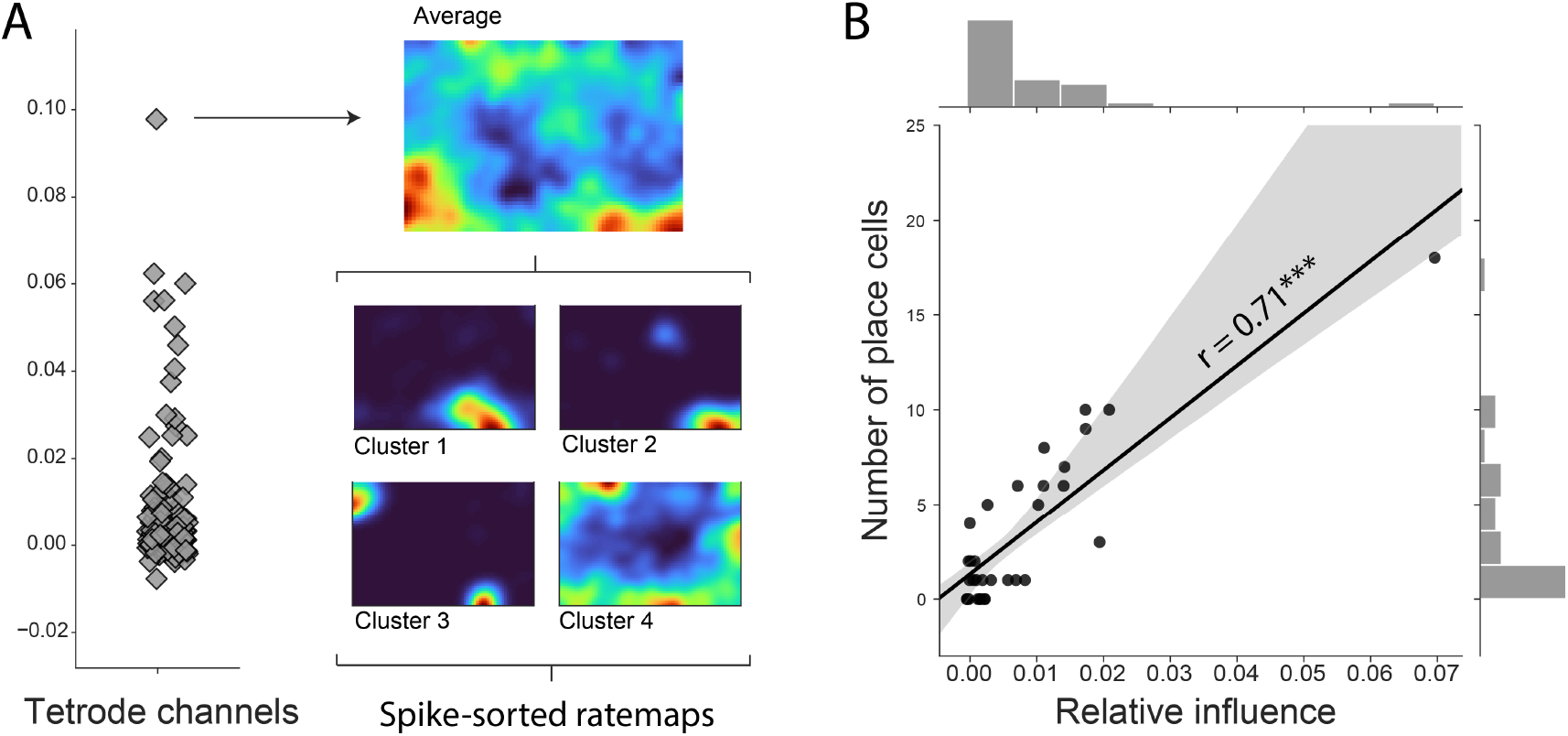
Influence of decoding across channels. (A) Channel influence scores for positional decoding (left). The most influential channel for the positional decoding has a high number of place cells (right). Average ratemap of all clusters (top, n=21) and four example clusters (bottom) shown. (B) influence scores per tetrode (average influence over 4 channels) highly correlates with number of place cells on the tetrode, indicating that the network is correctly identifying place cells as the spatially most informative neural correlate.

**Figure S4:**
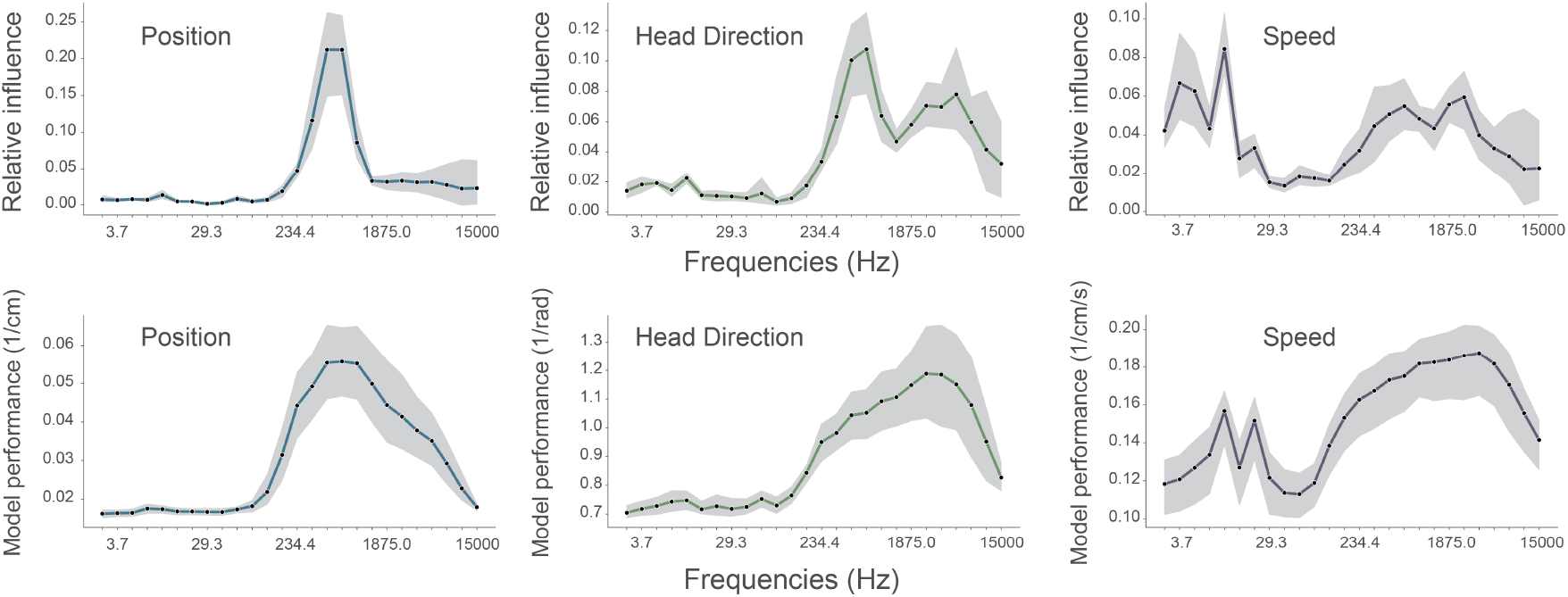
Performance of standard model compared with models trained on single frequency bands. To determine if the frequencies identified as important by our complete model matched those that were most informative on their own, we compared the influence plots (top row, same as Figure 3A in manuscript) generated for the standard model with accuracy plots from models trained on individual frequency bands (bottom row). In all cases 128 channel recordings from rodent CA1 were used to decode position (left column), head direction (middle column), and speed (right column). influence plots were constructed as before. Accuracy (bottom row) is simply defined as 1/decoding error and is not normalised relative to chance or ceiling performance, values were generated using the same convolutional neural network while only providing a single frequency band for training and testing. Although influence is not expected to be a simple linear function of accuracy, the results from the two methods were highly correlated: position, Spearman’s Rho=0.88 (p<0.001); head direction, Rho=0.82 (p<0.001); running speed, Rho=0.47 (p=0.02). For each frequency band we show the average cross-validation performance across five animals for three different behaviours and loss functions. Data from all animals, the shaded area indicates the 95% confidence interval.

**Figure S5:**
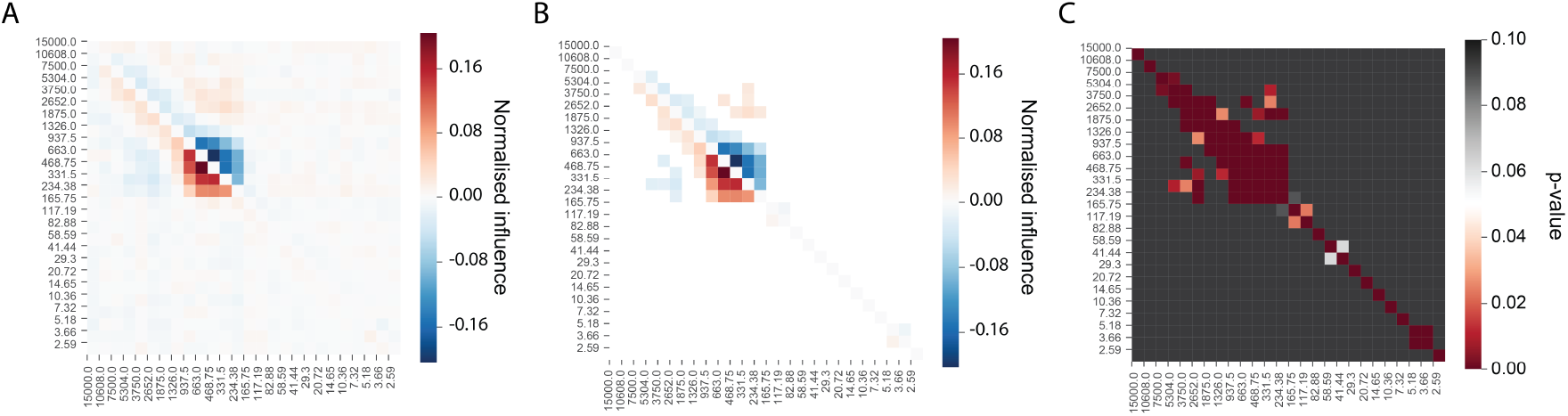
A subset of frequency pairs exhibit greater than expected decoding influence. We evaluated 325 frequency pairs to investigate the relative influence of conjoint frequencies versus the sum of their individual influence on the decoding of position. (A) For each frequency pair the combined influence - calculated by shuffling both together (*f_xy_*) - is compared to the summed influence of each alone (*f_x_* + *f_y_*). Lower triangle shows *f_xy_* − (*f_x_* + *f_y_*), upper triangle shows (*f_x_* + *f_y_*) − *f_xy_*. Positive, red, entries in lower triangular indicate frequency pairs with a combined influence greater than the sum of their individual influences. (B) Same a left matrix with non-significant entries removed. (C) P values for the data in the left matrix determined using the Wilcoxon signed rank test, Holm-Sidak corrections were applied for n=325 comparisons. There was a limited subset of frequency pairings in which the combinatorial influence on position decoding significantly exceeded the sum of individual frequencies - these were focused on the bands associated with place cell action potential (331.5Hz to 937.5Hz) and to a lesser extent on the ones associated with putative interneurons

**Figure S6:**
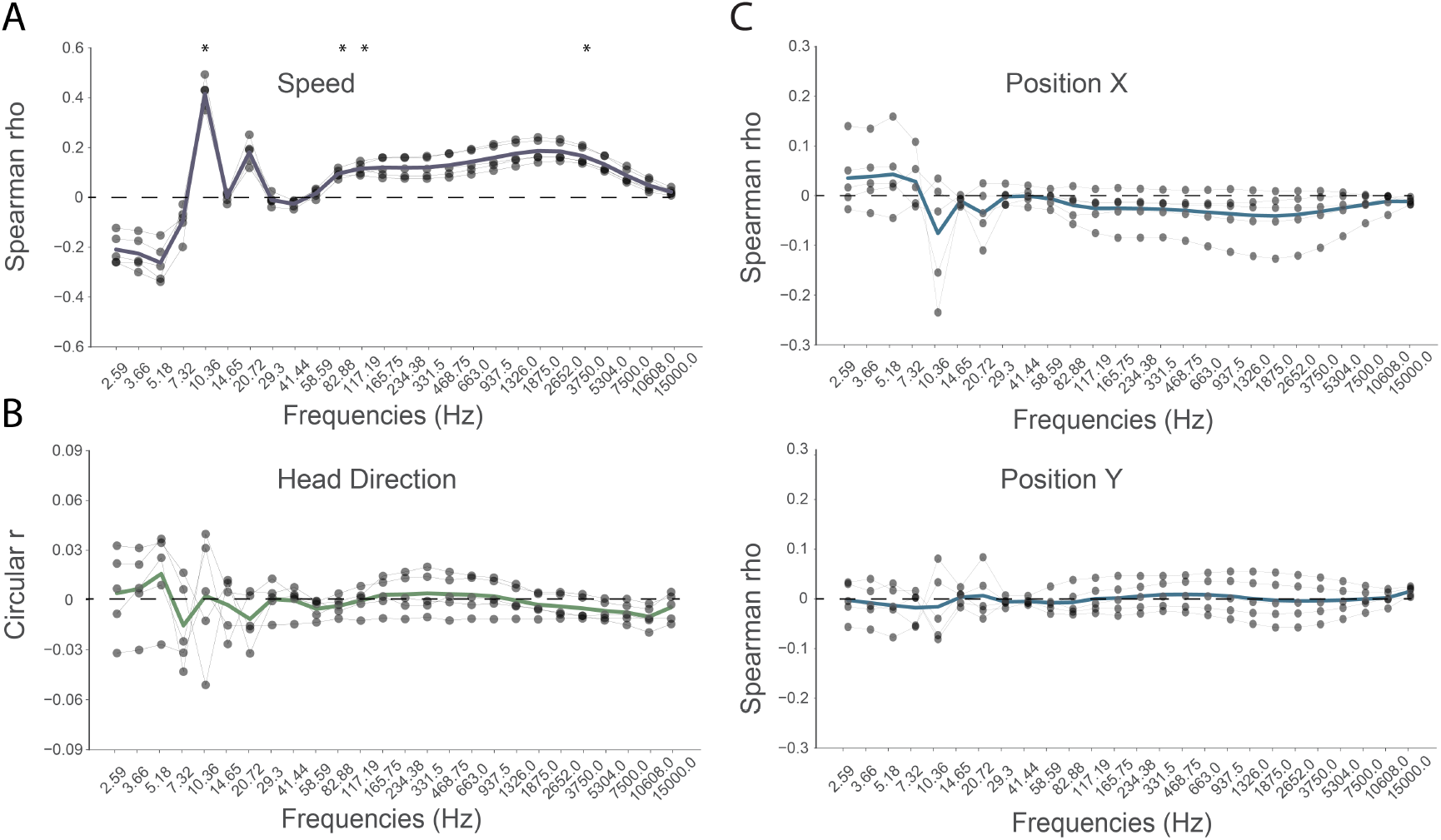
Running speed linearly correlates with power in multiple frequency bands. To identify simple relationships between behavioural variables and frequency bands we calculated the Spearman Rank Order Correlation Coefficient (speed and position) and Circular Correlation (head direction) between the wavelet transformed electrophysiological signal and the position, head direction and speed of the animal. (A) In the case of running speed, multiple frequency bands exhibited moderate correlations. In particular, as previously reported, the strongest relationship was present in the theta-band (10.36Hz, rho = 0.415), a relationship that was present in all five rats (p < 0.01). (B,C) In contrast no such relationship was found for the other spatial variables. Indeed, the strongest correlation identified was a negative relationship between x-axis position and power in the theta-band (10.36Hz, rho=−0.0759, but which was not significant, p = 0.9803). Grey points indicate correlation for each animal (n=5), bold line shows the mean of those.

**Figure S7:**
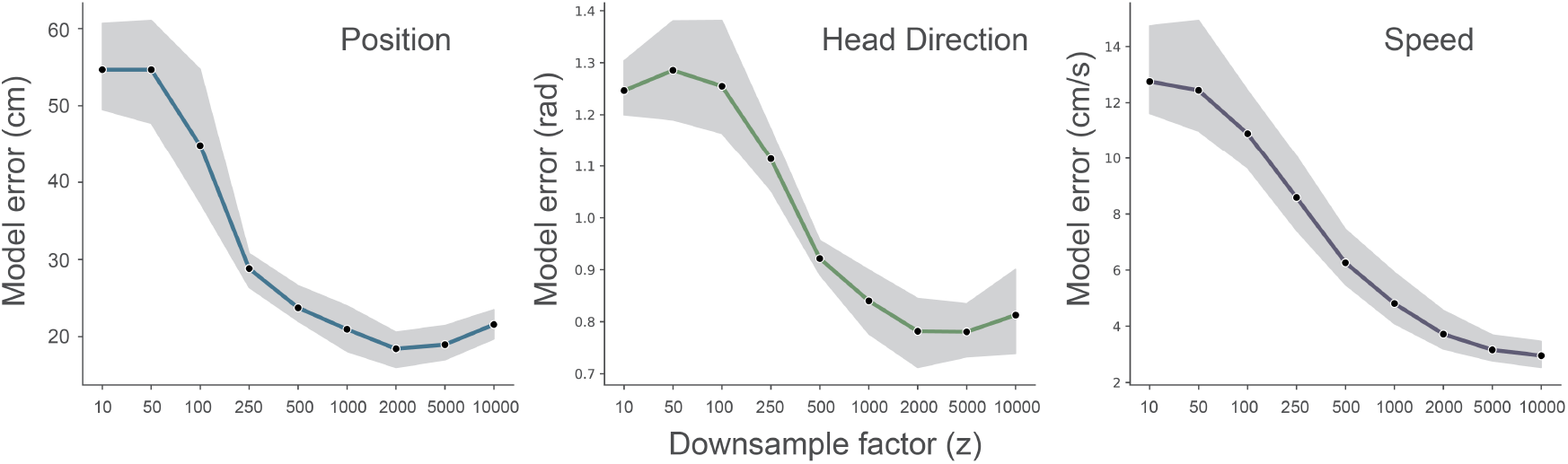
Effect of downsampling on model performance. We ran fully cross-validated experiments for different downsampling values of the wavelet transformed electrophysiological signal (z = [10…10000]). Data from one animal (R2478), the shaded area indicates the 95% confidence interval.

**Table S1:** Layer by layer architecture of the convolutional model. Note that the first layers 1-8 share the weights over the channel dimension while layers 9-15 share the weights across the time dimension. Layers 9 to 15 depict the kernel sizes and strides for the tetrode recordings with 128 channels. For recordings with different number of channels we adjust the number of downsampling layers to match the dimension of layer 15. Order of dimensions: Time, Frequency, Channels.

| Layer | Layer Type | Input Dimension | Kernel Size | Strides | Filters |
| --- | --- | --- | --- | --- | --- |
| 1 | 2D - Convolution | (64, 26, 128) | (3,3) | (2,1) | 64 |
| 2 | 2D - Convolution | (32, 26, 128) | (3,3) | (1,2) | 64 |
| 3 | 2D - Convolution | (32, 13, 128) | (3,3) | (2,1) | 64 |
| 4 | 2D - Convolution | (16, 13, 128) | (3,3) | (1,2) | 64 |
| 5 | 2D - Convolution | (16, 7, 128) | (3,3) | (2,1) | 64 |
| 6 | 2D - Convolution | (8, 7, 128) | (3,3) | (1,2) | 64 |
| 7 | 2D - Convolution | (8, 4, 128) | (3,3) | (2,1) | 64 |
| 8 | 2D - Convolution | (4, 4, 128) | (3,3) | (1,2) | 64 |
| 9 | 2D - Convolution | (4, 2, 128) | (3,2) | (1,2) | 64 |
| 10 | 2D - Convolution | (4, 2, 64) | (3,2) | (1,2) | 128 |
| 11 | 2D - Convolution | (4, 2, 32) | (3,2) | (1,2) | 128 |
| 12 | 2D - Convolution | (4, 2, 16) | (3,2) | (1,2) | 128 |
| 13 | 2D - Convolution | (4, 2, 8) | (3,2) | (1,2) | 128 |
| 14 | 2D - Convolution | (4, 2, 4) | (3,2) | (1,2) | 128 |
| 15 | 2D - Convolution | (4, 2, 2) | (3,2) | (1,2) | 128 |
| 16 | Fully Connected | (4, 512) | - | - | 1024 |
| 17 | Fully Connected | (4, ) | - | - | 2/1 |

